# Mechanical loading rescues mechanoresponsiveness in a human osteoarthritis explant model despite Wnt activation

**DOI:** 10.1101/2023.11.27.568718

**Authors:** R Castro-Viñuelas, N Viudes-Sarrión, AV Rojo-García, S Monteagudo, RJ Lories, I Jonkers

**Affiliations:** Department of Movement Sciences, Human Movement Biomechanics Research Group, KU Leuven, Leuven, Belgium; Department of Development and Regeneration, Skeletal Biology and Engineering Research Centre, Laboratory of Tissue Homeostasis and Disease, KU Leuven, Leuven, Belgium. Electronic address; Department of Development and Regeneration, Skeletal Biology and Engineering Research Centre, Laboratory of Tissue Homeostasis and Disease, KU Leuven, Leuven, Belgium. Electronic address; Department of Development and Regeneration, Skeletal Biology and Engineering Research Centre, Laboratory of Tissue Homeostasis and Disease, KU Leuven, Leuven, Belgium; Division of Rheumatology, University Hospitals Leuven, Leuven, Belgium. Electronic address; Department of Movement Sciences, Human Movement Biomechanics Research Group, KU Leuven, Leuven, Belgium; Skeletal Biology and Engineering Research Centre, KU Leuven, Leuven, Belgium. Electronic address

## Abstract

**Objectives:** Optimizing rehabilitation strategies for osteoarthritis necessitates a comprehensive understanding of chondrocytes’ mechanoresponse in both health and disease, especially in the context of the interplay between loading and key pathways involved in osteoarthritis development, like canonical Wnt signaling. This study aims to elucidate the role of Wnt signaling in the mechanoresponsiveness of healthy and osteoarthritic human cartilage.

**Methods:** We used an ex-vivo model involving short-term physiological mechanical loading of human cartilage explants. First, the loading protocol for subsequent experiments was determined. Next, loading was applied to non-OA explants with or without Wnt activation with CHIR99021. Molecular read-outs of anabolic, pericellular matrix and matrix remodeling markers were used to assess the effect of Wnt on cartilage mechanoresponse. Finally, the same set-up was used to study the effect of loading in cartilage from patients with established OA.

**Results:** Our results confirm that physiological loading maintains expression of anabolic genes in non-OA cartilage, but indicate a deleterious effect of Wnt activation in the chondrogenic mechanoresponsiveness. This suggests that loading-induced regulation of cartilage markers occurs downstream of canonical Wnt signaling. Interestingly, our study highlighted contrasting mechanoresponsiveness in the model of Wnt activation and the established OA samples, with established OA cartilage maintaining its mechanoresponsiveness, and mechanical loading rescuing the chondrogenic phenotype.

**Conclusion:** This study provides insights into the mechanoresponsiveness of human cartilage in both non-OA and OA conditions. These findings hold the potential to contribute to the development of strategies that optimize the effect of dynamic compression by correcting OA pathological cell signaling.

## 1. Introduction

Articular cartilage is a highly specialized type of connective tissue that covers the ends of the bones, providing a smooth, almost frictionless surface for articulation and facilitating the load transmission to the underlying subchondral bone^1,2^. During joint movement, articular cartilage is simultaneously exposed to axial compressive loading, tensile strain and shear forces^3,4^. Articular chondrocytes, the unique cell population within the joint cartilage, are highly mechanosensitive cells. Through mechanotransduction, they can convert externally applied mechanical stimuli conveyed by the extracellular and pericellular matrix (ECM and PCM), into molecular responses which in turn promote tissue adaptation^5,6^. Hence, mechanical signals are key factors that contribute to joint homeostasis but they are also implicated in the development of osteoarthritis (OA), the most common chronic joint disease affecting more than 500 million people globally^7^. OA is a disease of the whole joint but the loss of articular cartilage is its hallmark, with simultaneous pathological alterations in the subchondral bone, synovium and meniscus. In OA cartilage, the balance between anabolic and catabolic responses of the chondrocytes is distorted, dysregulating joint homeostasis and triggering structural damage to the articular cartilage.

The detailed molecular mechanisms of OA initiation and progression are not yet fully understood and, to date, no interventions are available to restore damaged cartilage or to consistently slow down disease progression^8^. A better understanding of the chondrocytes’ mechanoresponse in homeostasis and disease may not only help to better position drug interventions in the disease course but could also be used to develop mechanical loading strategies that are optimally tuned to synergize with the effect of drugs on the joint. This requires untangling the - likely complex - interactions of mechanical loading with different intracellular pathways that either promote or dysregulate cartilage homeostasis.

Increased activation of canonical Wnt signaling has been implicated in the development of OA through genetic studies and experimental evidence^9–11^. Wingless-related integration site (Wnt) signaling refers to a family of at least 19 different secreted proteins in humans that affect a large variety of biological processes. Upon binding with its receptors, signaling is initiated through different mechanisms, of which the β-catenin-dependent or canonical pathway is the best understood^12^. Wnt signaling has been identified as a pathway triggered upon injurious mechanical stimulation, hence contributing to post-traumatic OA development^13^. However, interactions between Wnt signaling pathway activation and the chondrocytes’ mechanoresponse in health and disease remain largely unexplored. In this study, we aim to study the role of Wnt signaling in the mechanoresponsiveness of healthy and osteoarthritic cartilage, upon short-term physiologic mechanical loading. We hypothesized that activation of Wnt signaling seen in OA will hamper the chondrogenic mechanoresponsiveness.

## 2. Materials and methods

### 2.1 Articular cartilage explants collection and culture

Informed consent and ethical approval of the University Hospitals Leuven Ethics Committee and Biobank Committee (Leuven, Belgium) (S56271) was obtained before performing this study. Non-OA human cartilage explants were harvested from the hips of 11 donors with no history of OA undergoing hip replacement surgery for osteoporotic or malignancy-associated fractures (8 women, 3 men, age 76.72 ± 10.88). OA human cartilage explants were obtained from the hips of 12 patients undergoing hip replacement surgery due to OA (8 women, 4 men, age 69.75 ± 6.79, supplementary table 1). First, cartilage was dissected into fragments using a scalpel and then rinsed in PBS supplemented with 1% (vol/vol) antibiotic/antimycotic (Gibco). The cartilage fragments were subsequently cut with an 8 mm diameter disposable biopsy punch (Robbins Instruments) and cultured for 24h in DMEM/F12 (Gibco), 10% fetal bovine serum (FBS, Gibco), 1% (vol/vol) antibiotic/antimycotic (Gibco), and 1% L-glutamine (Gibco) at 37°C and 5% CO_2_ humidified atmosphere before starting the experiments.

### 2.2 Wnt activation model

Articular cartilage explants from the non-OA donors were treated with 3 μM of Wnt agonist CHIR99021 (Sigma-Aldrich) or with 1 μL/mL of its vehicle (V) dimethyl sulfoxide (DMSO) (control group) during 24 hours before performing mechanical stimulation in the bioreactor. This concentration was based on literature^14^ and on previous data of the laboratory^15^.

### 2.3 Cartilage explants viability assay

Cartilage explants viability before and after treatment with CHIR99021 was measured using the resazurin-based PrestoBlue™ Cell Viability Reagent kit (ThermoFisher) according to manufacturer’s instructions. Briefly, samples were incubated with 10% PrestoBlue™ solution in culture medium for 2 hours at 37°C. Absorbances at 570 and 600 nm were then measured using a BioVert™ Synergy HT plate reader.

### 2.4 Ex vivo mechanical stimulation

Ex vivo mechanical uni-axial stimulation was applied to articular cartilage explants in the ElectroForce® BioDynamic 5210 bioreactor (TA Instruments) with the accompanying WinTest® Software, inside an incubator at 37°C and with 5% of CO_2_ (Figure 1a). All elements of the loading chambers were sterilized with UV light before use. Cartilage explants were measured in height and positioned in the sample holder of the loading chamber with the upper layer of the cartilage facing up, keeping sterile conditions. The two surfaces of the sample holder were brought together until the cartilage explant was secured. Once closed, the loading chambers were filled with culture medium (DMEM/F12 + 10% FBS + 1% antibiotic/antimycotic + 1% L-glutamine) and placed in the bioreactor.

**Figure 1.**
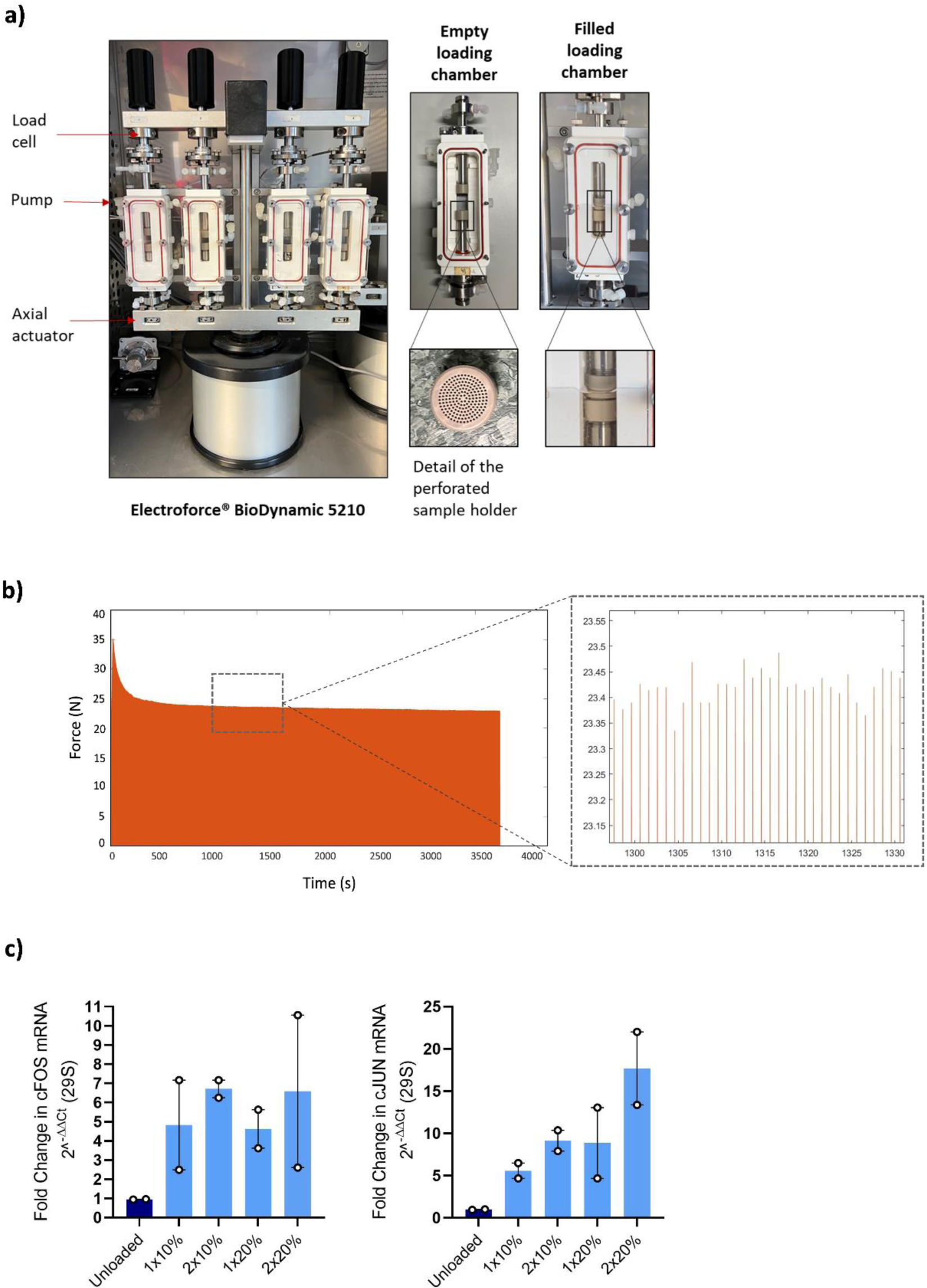
Ex vivo compressive loading experimental set up. **a)** The Electroforce® BioDynamic 5210 Bioreactor was used to apply cyclic compression to human cartilage explants by vertical movement of the axial actuator in a 5% CO_2_ incubator. Loading applied to the explants was recorded by the 225N load cell located on top of each loading chamber. Culture medium was added in sterile conditions through the pump until the loading chamber was filled and the perforated sample holder allowed for fluid flow during the loading protocol. **b)** Reaction force versus time curve recorded by the load sensor of the bioreactor, indicating an equilibrium in the reaction force after 300 seconds and zoom in version of the curve showing the applied cyclic loading. **c)** Testing of different loading protocols. Fold changes in mRNA levels of mechanosensitive genes cFOS and cJUN were evaluated under 4 different conditions. Data are expressed as individual data points representing each donor sample and mean ± SEM.

In a first experiment, the physiological loading protocol for use in the subsequent experiments was determined. To this end, four different protocols of compression with a sinusoidal waveform (Figure 1b) were tested: (**1**) 10% compressive strain (compression) at a frequency of 1Hz for 1 hour + 1 hour of free swelling, (**2**) 10% compression at 1Hz for 1 hour + 1 hour of free swelling + 10% compression at 1Hz for 1 hour + 1 hour of free swelling, (**3**) 20% compression at a frequency of 1Hz for 1 hour + 1 hour of free swelling, (**4**) 20% compression at 1Hz for 1 hour + 1 hour of free swelling + 20% compression at 1Hz for 1 hour + 1 hour of free swelling. Paired control explants were maintained under free swelling conditions at 37°C and 5% CO_2_.

Upon selection of the most relevant loading protocol, in a second experiment, mechanical loading was applied to non-OA explants in the presence and absence of Wnt activation with CHIR99021 to investigate the effect of Wnt activation on cartilage mechanoresponse. In this sense, four experimental conditions were tested for each non-OA donor: free swelling vehicle (control condition), load vehicle, free swelling + CHIR99021 and load + CHIR99021 (Figure 2a). In a final experiment, cartilage explants from patients with established OA were loaded and compared to free swelling explants (Figure 4a).

**Figure 2.**
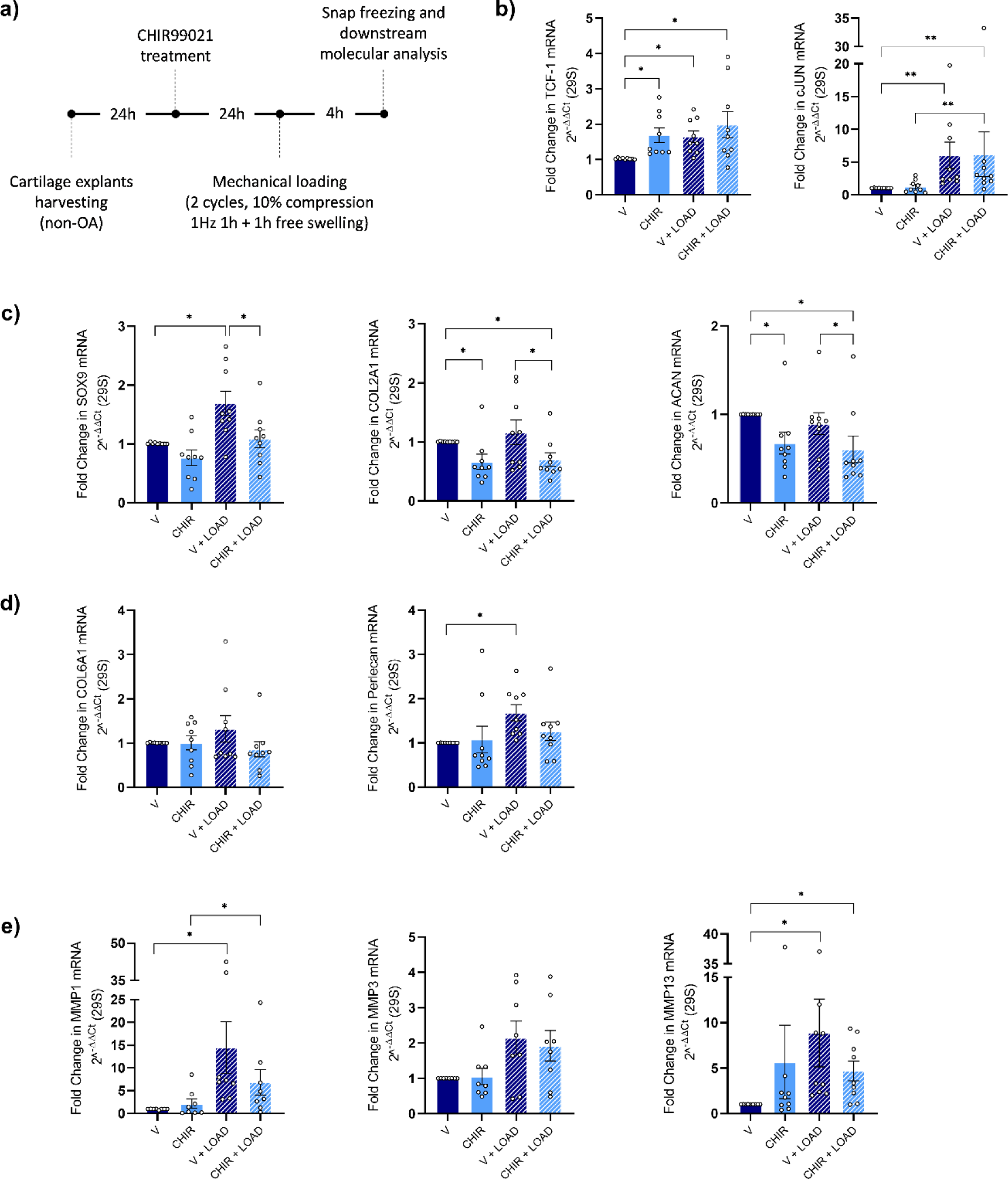
Activation of Wnt signaling alters cartilage mechanoresponsiveness. **a)** Schematic representation of the followed workflow. Real-time PCR analysis of **b)** positive control genes TCF1 and cJUN, **c)** marker of chondrogenesis SOX9, COL2A1 and ACAN, **d)** pericellular matrix marker genes COL6A1 and Perlecan and **e)** matrix remodeling enzymes MMP1, MMP3 and MMP13 in human non-OA articular cartilage explants after mechanical stimulation (LOAD) and/or treatment with 3μM CHIR99021 (CHIR) or vehicle (V). Data are expressed as individual data points representing each donor sample and mean ± SEM in all treatment conditions (n = 9, *P* < 0.05, Tukey corrected for 6 tests in linear mixed model).

### 2.5 RNA extraction of human cartilage explants

After completion of the loading protocol, loaded cartilage explants together with the paired controls were immediately washed with PBS + 1% antibiotic/antimycotic, snap-frozen in liquid nitrogen and stored at -80°C till RNA extraction was performed.

Total RNA extraction was performed following a modified protocol from Dell’accio et al 2008. Briefly, frozen explants were cut in small pieces of about 0.5 mm with a scalper while kept on ice and introduced in tissue grinding CKMix50-R 2 ml tubes (Bertin Technologies) containing 500 µl of TRIzol™ reagent (Invitrogen). Tubes were placed in a Precellys Evolution tissue homogenizer (Bertin Technologies) and cartilage tissue was homogenized following a protocol consisting in 3 cycles of 15 seconds at 8600 rpm at 4°C followed by 1 min of rest on ice. After homogenization, samples were kept on ice for 20 min and cellular debris was removed by centrifugation at 10000 g for 10 min at 4°C. Chloroform was added to the supernatant and tubes were shaken by hand for 30 seconds, then incubated on ice for 2 min and centrifuged at 10000 g for 15 min at 4°C. Aqueous phase was transferred to a fresh tube and an equal volume of TRIzol™ was added and phase separation was repeated. Aqueous phase was taken, mixed with equal volume of isopropanol and incubated at -20°C overnight. After overnight incubation, samples were centrifuged at 12000 g for 30 min at 4°C. Supernatant was poured-off, RNA pellet was washed with 1 ml of 70% ethanol and centrifuged for 5 min at 7000 g at 4°C. Supernatant was discarded and pellets were air dried for 5 min at room temperature before resuspended in RNAse-free water. Quantity and quality of the RNA was measured with a NanoDrop spectrophotometer (ThermoFisher Scientific).

### 2.6 Gene expression analysis

cDNA was synthesized using the Revert Aid H minus First Strand cDNA synthesis kit (Thermo Fisher Scientific) according to the manufacturers’ instructions. Relative quantity of mRNA was determined using StepOne Real-Time PCR Systems with the Maxima SYBRgreen qPCR master mix system (Thermo Fisher Scientific). Relative gene expression was calculated following normalization to housekeeping gene 29S mRNA levels using comparative Ct (cycle threshold) method. The following PCR conditions were used: incubation for 10 min at 95 °C followed by 40 amplification cycles of 15 s of denaturation at 95 °C 14 followed by 45 s of annealing-elongation at 60 °C. Melting curve analysis was performed to determine the specificity of the amplicons. Primers used in qPCR are listed in supplementary table 2.

### 2.7 Mechanical characterization of explants

Young’s modulus of the non-OA and OA cartilage explants was calculated from the slope of stress-strain curve during loading and after reaching equilibrium, based on the recorded force and displacement data (250 N, TA Instruments), using an in-house developed code (Matlab 2021b, Mathworks).

### 2.8 Statistical analysis

Data analysis and graphical presentation were performed with GraphPad Prism version 8 and R-Studio Version 1.1.463 (packages: car, rstatix, tidyverse, ggpubr, lme4, emmeans, MASS, multcomp, lmerTest, EnvStats, readr, effects, sjPlot, Matrix, lattice). Data are presented as mean ± SEM and as individual data points, as indicated in figure legends. Gene expression data were log-transformed for statistical analysis. Difference of means from raw data and CI corresponding with log transformed data used for statistical analysis are reported in the text. For comparisons between non-OA and OA groups, unpaired 2-tailed Student’s t test was used. For comparisons between 2 groups with non-independent data (different experimental conditions in samples from one donor), paired 2-tailed Student’s t test was used. A linear mixed model was used for the experiments with CHIR99021-treated samples, with individual donor as random factor, to study interactions and main effects between independent and categorical variables (R script section 13). Tukey correction was used for multiple comparisons. Data are reported by t values and exact P values in supplementary table 3 where applicable. P values of pairwise comparisons are indicated in the graphs. Model assumptions were further checked by QQ plots and homoscedasticity plots. Homogeneity of variance was evaluated by standardized residuals versus fit plot. P values of less than 0.05 were considered significant.

## 3. Results

### 3.1 Definition of a short-term physiologic loading protocol for human cartilage explants

We first set out to establish a short-term loading protocol that induces the expression of the well-established mechanosensitive genes cFOS and cJUN (Figure 1c). The four loading protocols increased the expression levels of cFOS compared to the unloaded control, but the protocol consisting in 2 cycles of 10% compression led to the highest upregulation. The expression levels of cJUN were also increased with all the tested loading protocols. In particular, the protocol consisting in 2 cycles of 20% compression (protocol 4) led to the highest upregulation compared to the unloaded control, followed by 2 cycles of 10% compression. Based on these results, and considering published evidence suggesting 20% compression as overloading^16,17^, the protocol consisting of 2 cycles of 10% compression for 1 hour (protocol 2) was selected as loading protocol for all subsequent experiments. Furthermore, given that cJUN expression levels showed a dose-dependent response, it was chosen as positive control gene for mechanical stimulation in the follow-up experiments.

### 3.2 Excessive Wnt signaling compromises the chondrogenic mechanoresponse

Next, we investigated to what extent excessive Wnt activation induced at baseline (prior to loading) could affect cartilage mechanoresponsiveness (Figure 2a-d). Explants were therefore treated with Wnt agonist CHIR99021. While viability of cells in non-OA human cartilage explants was not affected by the treatment with 3 µM CHIR99021 for 24h (Supplementary figure 1), upregulation of the main Wnt target gene TCF-1 was observed after treatment in all donors, compared to the vehicle condition (difference of means, 0.662 [95% CI (-0.369) – (-0.0174)], *P* = 0.0269). When compressive dynamic loading was applied, expression levels of TCF-1 also increased compared to the control condition, both in the presence and in the absence of CHIR99021, suggesting that TCF-1 is also mechanoresponsive (difference of means, 0.966 [95% CI (-0.407) – (-0.055)], *P* = 0.0064 and 0.618 [95% CI (-0.365) – (-0.0126)], *P* = 0.032, respectively). The levels of cJUN were evaluated as a positive control of mechanical stimulation: Expression levels of cJUN increased upon loading compared to the control condition both in the presence and in the absence of CHIR99021 (difference of means, 5.162 [95% CI (-0.908) – (-0.121)], *P* = 0.0065) and 5.011 [95% CI (-1.007) – (-0.221)], *P* = 0.001, respectively). No differences were observed between control and CHIR99021-treated samples (difference of means, 0.235 [95% CI (-0.37) – (0.416)], *P* = 0.9986), indicating similar activation of the mechanosensitive pathways even in the presence of excessive Wnt activation.

Mechanical loading alone upregulated SOX9 mRNA expression (difference of means, 0.683 [95% CI (-0.382) – (-0.013)], *P* = 0.0324), while its expression levels were significantly reduced when mechanical loading was applied in the presence of CHIR99021 compared to loading without CHIR99021 (difference of means, -0.601 [95% CI 0.014-0.384], *P* = 0.0307). COL2A1 mRNA expression did not show consistent modifications after loading (with upregulation being observed in samples from 5 non-OA donors but downregulation in samples from 4 non-OA donors). However, CHIR99021 treatment significantly reduced the expression levels of COL2A1 (difference of means, -0.339 [95% CI 0.039-0.416], *P* = 0.0131) and this decrease was not compensated by mechanical stimulation, being significantly lower than the loading condition (difference of means, -0.459 [95% CI 0.010-0.387], *P* = 0.0358). ACAN mRNA expression showed no changes upon mechanical stimulation compared to the control condition (difference of means, -0.106 [95% CI (-0.098) – (0.263)], *P* = 0.6047). Similar to the response of COL2A1, CHIR99021 treatment significantly reduced the expression levels of ACAN (difference of means -0.325 [95% CI 0.037-0.397], *P*= 0.0136) and this decrease was not rescued after loading. Dynamic compression in the presence of CHIR99021 significantly reduced the mRNA expression levels of ACAN compared to loading in the untreated samples (difference of means -0.291 [95% CI 0.032-0.393], *P* = 0.0161).

As the PCM is an important zone for the chondrocyte’s mechano-sensing and -response, we tested the effects of loading and Wnt activation on specific PCM markers. No changes were observed in the mRNA expression levels of COL6 in neither of the tested experimental conditions (difference of means, -0.004 for V vs CHIR [95% CI (-0.186) – (0.307)], *P* = 0.9074; 0.318 for V vs LOAD [95% CI (-0.293) – (0.199)], *P* = 0.9545; -0.147 for V vs CHIR LOAD [95% (-0.113) – (0.380)], *P*= 0.4665; -0.143 for CHIR vs CHIR LOAD [95% CI (0.174) – (0.319)], *P* = 0.8534; -0.465 for LOAD vs CHIR LOAD [95% CI (-0..66) –(0.426)], *P* = 0.2147), but dynamic compressive loading significantly upregulated Perlecan expression (difference of means, 0.681 [95% CI (-0.404) – (-0.006)], *P* = 0.0412). This loading-induced increase in Perlecan mRNA expression levels was lost in the presence of CHIR99021 (difference of means, 0.255 [95% CI (-0.254) – (0.144)], *P* = 0.8779).

Tissue remodeling enzymes such as MMPs are typically associated with cartilage catabolism. We also evaluated their expression levels upon loading and Wnt stimulation. Surprisingly, dynamic compressive loading upregulated MMP1 and MMP13 (difference of means, 13.48 for MMP1 [95% CI (-1.612) – (-0.326)], *P* = 0.0017; 7.855 for MMP13 [95% CI (-1.154) – (-0.302)], *P* = 0.0003), but less consistently MMP3 (difference of means, 1.143 [95% CI (-0.594) – (0.161], *P* = 0.4137). When mechanical loading was applied in combination with the CHIR99021 treatment, there was a downward trend in the mRNA expression levels of MMP1 and MMP13 in comparison with loading alone (difference of means, -7.64 for MMP1 [95% CI (-0.247) – (1.040)], *P* = 0.3501; -4.178 for MMP13 [95% CI (-0.242) – (0.609)], *P* = 0.6498).

In summary, our data suggest that loading-induced regulation of different cartilage anabolic and catabolic markers is impaired when Wnt signaling is activated. These results suggest that Wnt levels affect the complex loading-induced molecular changes, indicative of loading-induced regulation of cartilage markers occurring downstream of Wnt signaling.

### 3.3 Wnt signaling levels are increased in human explants with established OA

After investigating the mechano-responsiveness upon Wnt activation, we evaluated Wnt signaling levels in cartilage explants with established OA prior to loading: Baseline mRNA expression levels of two Wnt target genes, TCF-1 and LEF-1, were upregulated in OA compared to non-OA cartilage (difference of mean, 0.164 for TCF-1 [95% CI 0.146-1.19], *P* = 0.0152; 0.024 for LEF-1 [95% CI 0.005-0.043], *P* = 0.0153). Moreover, the mRNA expression levels of Wnt antagonist DKK1 were significantly lower in the cartilage of OA patients compared to non-OA donors (difference of mean, -0.338 [95% CI (-0.995) – (-0.007)], *P* = 0.0471) (Figure 3).

**Figure 3.**
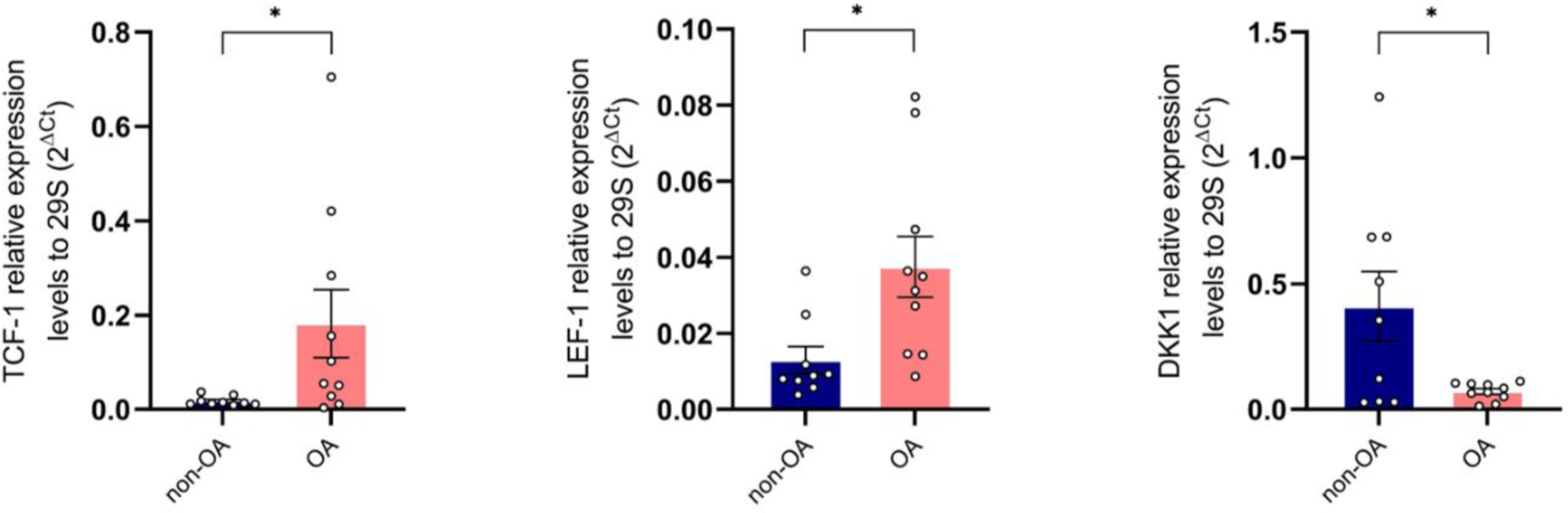
Human OA cartilage explants showed increased activation of the Wnt signaling cascade compared to non-OA cartilage explants. mRNA expression levels of the Wnt target genes TCF-1 and LEF-1 and the Wnt antagonist DKK1 in human non-OA (left) and OA (right) cartilage explants. n = 9, *P* < 0.05 by unpaired t-test.

### 3.4 Established OA cartilage preserves its mechanoresponsiveness and short-term physiologic mechanical loading rescues the chondrogenic phenotype

To investigate whether human cartilage explants with established OA show impaired mechanoresponsiveness, we applied the same mechanical loading protocol as above to explants harvested from 9 OA patients. Our first observation was that OA cartilage explants in the absence of loading show lower levels of the chondrogenic markers (SOX9, COL2A1 and ACAN), and the PCM-related marker COL6A1 than the non-OA samples (difference of mean, 4.778 for SOX9 [95% CI 0(-0.908) – (-0.299)], *P* = 0.0007; 6.099 for COL2A1 [95% CI (-0.979) – (0.0101)], *P* = 0.1046; 4.018 for ACAN [95% CI (-1.037) – (-0.224)], *P* = 0.0046; 1.118 for COL6A1 [95% CI (-1.283) – (-0.737)], *P* <0.0001) (Supplementary figure 2).

Although we hypothesized that human cartilage explants with established OA would show an impaired response to short-term mechanical stimulation, we observed that explants from OA patients positively responded to dynamic cyclic compression (Figure 5b). cJUN mRNA expression levels were significantly upregulated upon mechanical stimulation in all samples (difference of mean, 0.3 [95% CI, 0.357-1.151], *P* = 0.0024), showing that cartilage with established OA is still mechanoresponsive. In contrast to the CHIR99021-treated explants, which showed a reduced chondrogenic mechanoresponse, dynamic compression of OA cartilage explants led to significant increase in the mRNA expression levels of SOX9, COL2A1 and ACAN (difference of mean, 0.351 for SOX9 [95% CI, 0.289-0.925], *P* = 0.0023; 3.259 for COL2A1 [95% CI, 0.333-1.003], *P* = 0.0018; 4.702 for ACAN [95% CI, 0.016–0.593], *P* = 0.0408) partially rescuing the “OA phenotype”. Interestingly, the effect size (fold change to unloaded controls) was significantly higher in the OA samples compared to the non-OA cartilage explants. Given that no significant differences in overall tissue stiffness (Supplementary figure 4) were observed between OA and non-OA explants, indicative of similar intra-tissue strains being applied, this increased anabolic response suggests that cartilage from patients with established OA may be more receptive to mechanical stimulation than non-OA cartilage (Supplementary figure 3).

**Figure 4.**
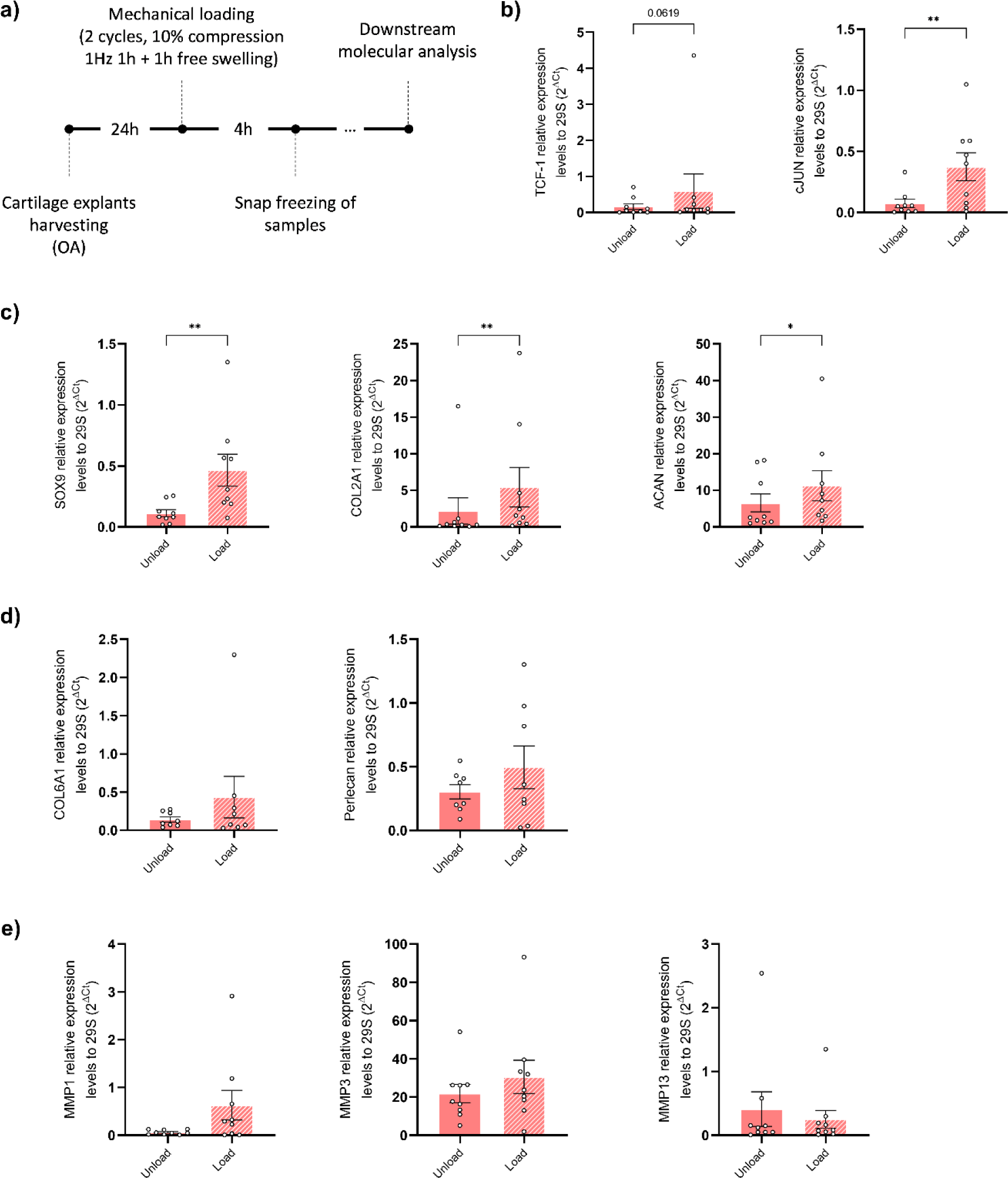
Effect of compressive mechanical loading on the gene expression profile of OA articular cartilage explants. **a)** Schematic representation of the followed workflow. Real-time PCR analysis of **b)** positive control genes TCF-1 and cJUN, **c)** marker of chondrogenesis SOX9, COL2A1 and ACAN, **d)** pericellular matrix marker genes COL6A1 and Perlecan and **e)** matrix remodeling enzymes MMP1, MMP3 and MMP13 in unloaded and loaded OA cartilage samples. Data are expressed as color-coded individual data points, representing each donor sample. n = 9, *P* < 0.05 by paired t-test.

On the other hand, the pattern of mechanoresponse of the PCM-related markers COL6A1 and Perlecan, and the catabolic enzymes MMP1, MMP3 and MMP13 was not consistent, observing upregulation upon mechanical loading in some samples but downregulation in other samples. To note, the magnitude of the response to dynamic compression in terms of MMP13 expression was reduced in OA compared to non-OA cartilage samples (difference of means, -7.536 [95% CI, (-1.208) – (0.3586)], *P* = 0.0012), suggesting a potential role for mechanical loading in regulating cartilage matrix remodeling.

## 4. Discussion

Understanding the molecular responses of cartilage to mechanical loading is key to the development of regenerative strategies that use mechanical signals to control cellular responses in an attempt to restore cartilage homeostasis and halt or slow down structural cartilage degradation. In this context, it is important to investigate potential interactions of OA-disease-specific signaling pathways that may offset chondrocytes’ mechanoresponsiveness in relevant *in vitro* models, that represent the *in vivo* human micro-environment. Here, we used human cartilage explants from both non-OA donors and OA patients, with the purpose of investigating the effect of short-term physiological mechanical loading in the presence of Wnt activation, as seen in human cartilage samples with established OA at baseline.

Upon activation of Wnt signaling in non-OA cartilage, the positive effect of cyclic compressive loading on the chondrogenic mechanoresponse was blunted. This effect was especially evident in the gene expression levels of SOX9, with CHIR99021 treatment altering the loading-induced increase of SOX9 mRNA levels. Interestingly, compressive loading was not able to maintain the expression levels of COL2A1 and ACAN in the presence of CHIR99021, suggesting a negative impact of Wnt activation to chondrogenic mechanoresponsiveness. These results are in line with previous observations in the tissue engineering field^18^, where high β-catenin levels suppressed the enhanced SOX9 protein and gene expression levels and GAG-stimulation after loading, whereas COL2A1 and ACAN gene expression levels were observed to be less load sensitive. In our study, similar trends were also observed for the PCM markers COL6A1 and Perlecan. Cyclic compressive loading in the absence of CHIR99021 significantly increased Perlecan levels, which is in accordance with a previous study in bovine chondrocytes^19^. Conversely, we observed a reduced response to loading after Wnt activation for COL6A1 and Perlecan, albeit not significant, probably due to interindividual variability in the human explant samples. Interestingly, the selected loading protocol led to upregulation of MMP1, MMP3 and MMP13 in the non-OA explants. Similar to our study, Blain *et al.* observed increased expression of MMPs in bovine cartilage explants upon loading^20^. Since cartilage mechanotransduction involves ECM adaptation to mechanical cues^6^, loading-induced MMPs activity may be required to promote adaptive remodeling of cartilage tissue. In fact, research has suggested that MMPs are important regulators of homeostatic tissue remodeling^21,22^. The mechanosensitive genes cFOS and cJUN are known to play a role in the transcriptional regulation of MMPs^23^, which can in addition explain the increased expression of the MMP genes in the loading condition. In our experiments, Wnt activation reduced the loading-induced upregulation of MMP1 and MMP13 despite unaltered cJUN expression, further suggesting the need of MMP1 and MMP13 to support loading-induced tissue remodeling.

Altogether, these results confirm that physiological loading maintains expression of anabolic, chondrogenic genes in non-OA human articular cartilage, which contributes to cartilage homeostasis, and suggests a deleterious effect of Wnt signaling activation on the chondrogenic mechanoresponsiveness. Targeted inhibition of Wnt signaling in OA may therefore boost the beneficial effect of compressive loading to promote cartilage anabolism. However, it is well known that the OA molecular fingerprint has a complex nature, affecting multiple and often overlapping signaling pathways^24^.

Indeed, mechanical loading rescued the chondrogenic mechanoresponsiveness in the established OA-explant model despite Wnt activation. Dynamic compressive loading led to a significant increase in the expression levels of the chondrogenic markers SOX9, COL2A1 and ACAN, in the presence of higher Wnt signaling levels than in non-OA cartilage. Given that no significant differences were observed in terms of tissue stiffness this finding suggests that established OA cartilage was more susceptible to mechanical stimulation than cartilage from non-OA donors as intra-tissue strain was comparable. These results are in contrast with other works^25^ who reported a decrease in COL2A1 and ACAN, and no changes in SOX9 following long term mechanical stimulation (3 days) in OA cartilage. As in the mentioned study no comparison with non-OA samples was performed, it is difficult to differentiate OA-related factors from factors related to tissue adaptation following the long-term culture. In this respect, the results of Dolzani and colleagues suggests impaired mechanoadaptation of OA cartilage, while our study suggests intact immediate mechanoresponsiveness. This observation is in line with previous studies^26,27^ that report an initial compensatory higher synthesis of COL2A1 and ACAN in OA samples. In fact, our observation that MMP13 is less responsive to loading in the established OA explants could be an indicative of potential impaired long-term adaptation. Future studies applying long-term mechanical loading are needed to gain further insights into the mechanoadaptive capacities of the tissue in the presence of the OA pathology and to clarify if and how the intact mechanoresponsiveness to compressive loading in OA cartilage can be maintained over time.

Interestingly, our study highlighted obvious differences in mechanoresponse in the model of Wnt activation as compared to the established OA samples. These discrepancies could be explained by the existence of OA-related mechanisms that antagonize with the canonical Wnt pathway, like noncanonical Wnt5a ligands^28^ or the noncanonical Ca2+/CaMKII-dependent pathway^29^, known to reciprocally inhibit canonical Wnt signaling. From this perspective, future studies will need to elucidate whether activation of noncanonical Wnt pathway or inhibition of the β-catenin-dependent pathway affect the cellular response to mechanical stimulation. Finally, given the difference in duration of Wnt signaling activation between the established OA-explants and the model with CHIR99021, the observed differences need to be further explored in the context of the differential impact of Wnt signaling on cartilage mechanoresponsiveness in early *vs* established OA. Considering that physical therapies are one of the first lines of treatment for OA^30,31^, this may have consequences when it comes to defining staged and personalized treatment strategies for OA patients. It is possible that at the early stages of the disease, when molecular derangement occurs^32^, physiotherapy treatments should be provided in combination with Wnt modulators, whereas short-term dynamic loading regimes could be used in more advanced stages to stimulate cartilage anabolism.

## 5. Conclusions

In conclusion, this study sheds light into human articular cartilage mechanoresponsiveness both in non-OA and OA conditions. We show that dynamic compressive loading contributes to cartilage homeostasis even during OA, but targeted early activation of canonical Wnt signaling compromises the chondrogenic mechanoresponse. This knowledge has the potential to contribute to the development of strategies that optimize the clinical effect of dynamic compression by correcting/optimizing OA pathological cell signaling.

## 6. Acknowledgements

We are grateful to the UZ Leuven nursing staff for their efforts to provide cartilage samples for this work.

## 7. Author contributions

R. Castro-Viñuelas: experimental design; acquisition, analysis and interpretation of data; drafting the manuscript; final approval of the version to be published; agreement to be accountable for all aspects of the work.

N. Viudes-Sarrión: acquisition, analysis and interpretation of data; drafting the manuscript; final approval of the version to be published; agreement to be accountable for all aspects of the work.

AV. Rojo-García: analysis of data for the work; final approval of the version to be published; agreement to be accountable for all aspects of the work.

S. Monteagudo: interpretation of the data; critical reviewing for important intellectual content; final approval of the version to be published; agreement to be accountable for all aspects of the work.

RJ Lories: conception of the work; interpretation of the data; critical reviewing for important intellectual content; final approval of the version to be published; agreement to be accountable for all aspects of the work.

I Jonkers: conception of the work; interpretation of the data; critical reviewing for important intellectual content; final approval of the version to be published; agreement to be accountable for all aspects of the work.

## 11. Figures and figure legends

**Supplementary figure 1.**
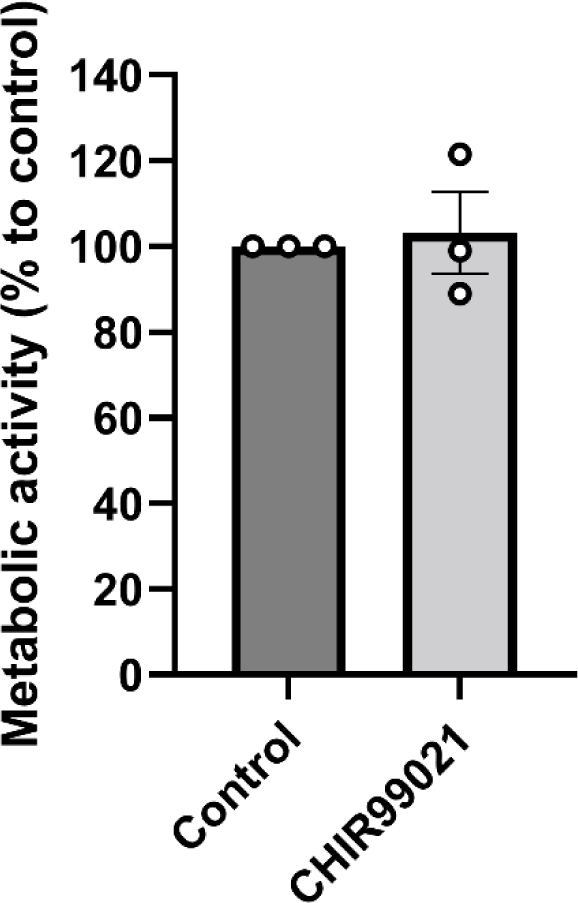
Cartilage explant viability. Each individual data pointt represents the mean of three technical replicates. Data are expressed as mean ± SEM of the metabolic activity relative to the untreated control (n = 3).

**Supplementary figure 2.**
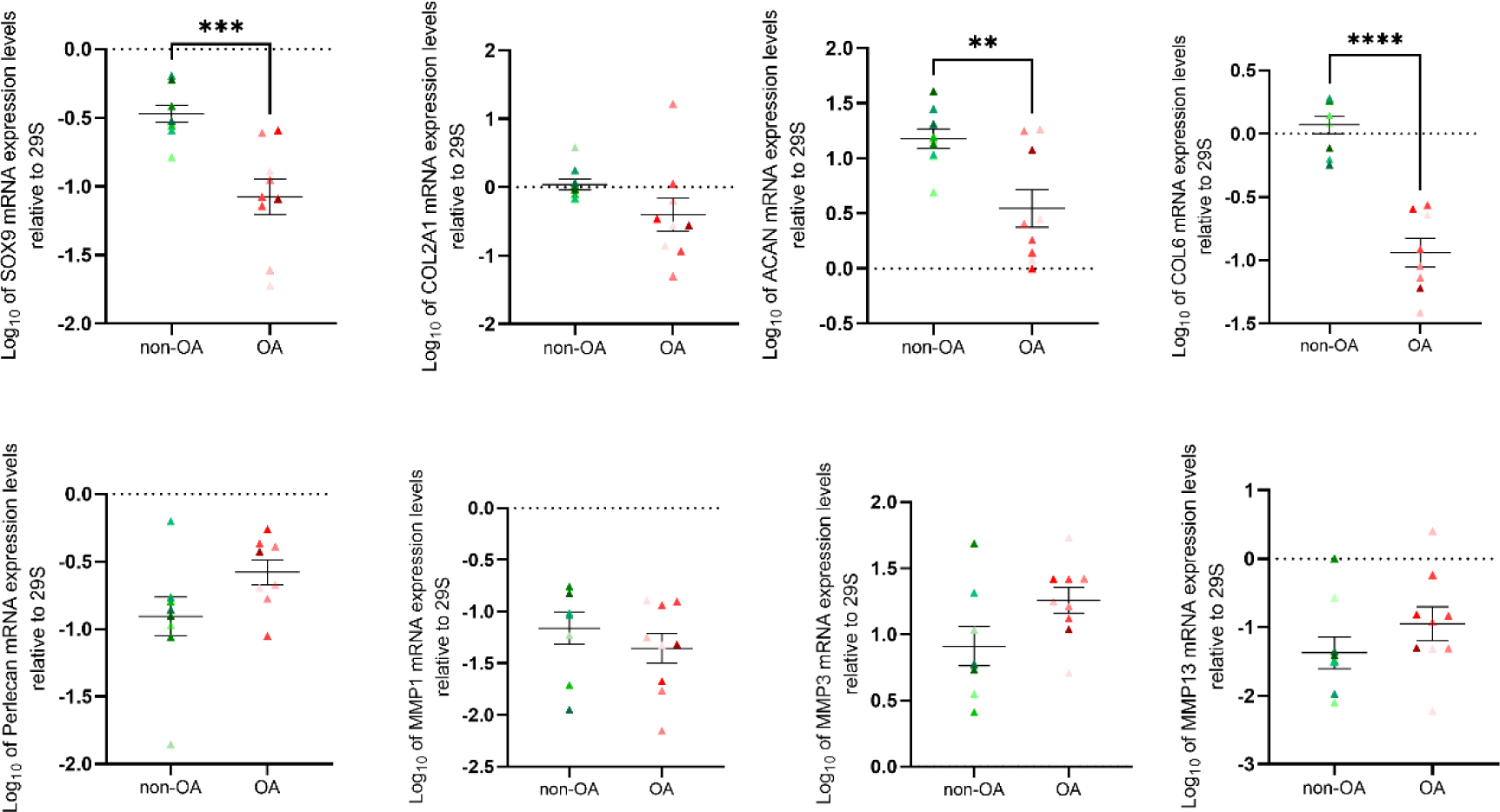
Basal levels of anabolic chondrogenic, pericellular matrix and matrix remodeling markers are lower in OA cartilage than non-OA cartilage. SOX9, COL2A1, ACAN, COL6A1, Perlecan, MMP1, MMP3 and MMP13 mRNA expression in unloaded non-OA and OA cartilage samples by qRT-PCR. Data are expressed as color-coded individual data points, representing each donor sample and mean ± SEM (log_10_ of mRNA relative expression levels). n = 9 per group, *P* < 0.05 by unpaired t-test. Data in the graph come from experiments of figure 2 and figure 4.

**Supplementary figure 3.**
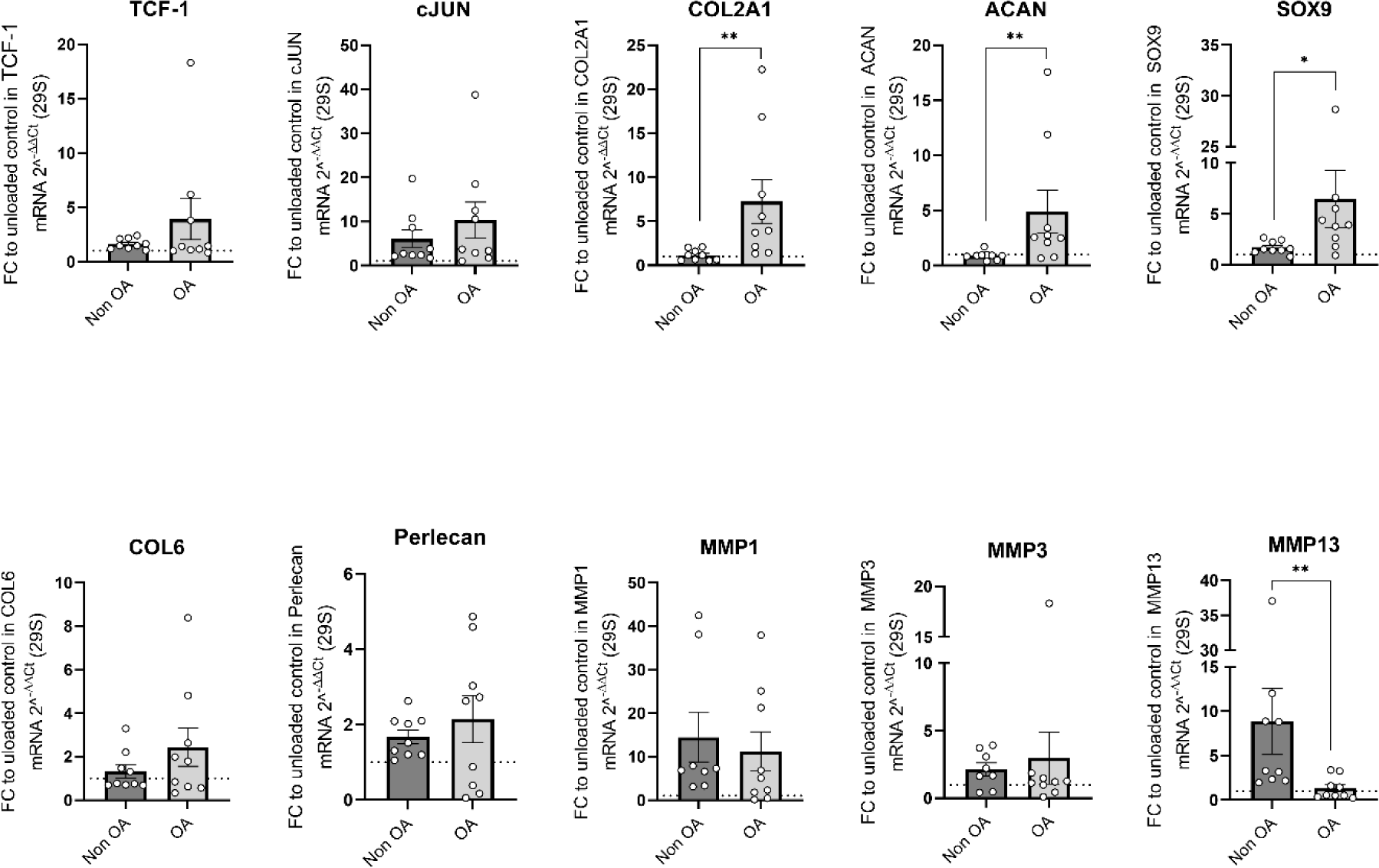
OA cartilage responds more to loading than non-OA cartilage. TCF-1, cJUN, SOX9, COL2A1, ACAN, COL6A1, Perlecan, MMP1, MMP3 and MMP13 mRNA expression in loaded samples from non-OA and OA cartilage by qRT-PCR. Data are expressed as individual data points, representing each donor sample and as mean ± SEM [fold change (FC) to unloaded control]. N = 9 per group, *P* < 0.05 by unpaired t-test. Data in the graph come from experiments of figure 2 and figure 4.

**Supplementary figure 4.**
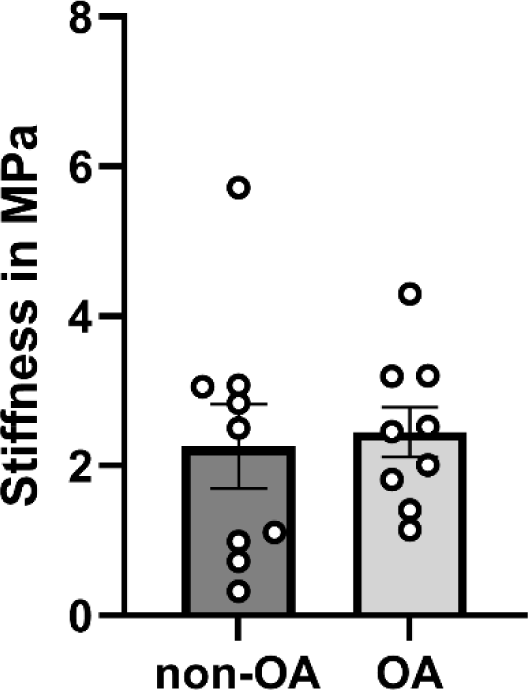
Mechanical characterization of human non-OA and OA cartilage explants. Young’s modulus of the non-OA and OA cartilage explants was calculated with the slope of stress-strain curve during loading in the bioreactor. n = 9 per group, *P* < 0.05 by unpaired t-test.

## 12. Supplementary tables

**Supplementary Table 1.** Patients’ data.

| Type | Age (years) | Gender | Experiment |
| --- | --- | --- | --- |
| non-OA | 85 | Female | Establishment of the physiologic loading protocol (figure 1) |
|  | 44 | Female |  |
|  | 77 | Female | Effect of Wnt signaling in cartilage mechanoresponsiveness (figure 2 and 3, supplementary figures 2, 3 and 4) |
|  | 80 | Female |  |
|  | 82 | Male |  |
|  | 74 | Male |  |
|  | 84 | Female |  |
|  | 80 | Female |  |
|  | 80 | Female |  |
|  | 83 | Female |  |
|  | 75 | Male |  |
| OA | 57 | Male | Presto Blue assay (supplementary figure 1) |
|  | 68 | Female |  |
|  | 76 | Male |  |
|  | 69 | Female | Wnt target genes levels and effect of physiologic mechanical loading in in OA cartilage mechanoresponse (figure 3 and 4, supplementary figures 2, 3 and 4) |
|  | 69 | Female |  |
|  | 71 | Female |  |
|  | 75 | Female |  |
|  | 84 | Female |  |
|  | 62 | Female |  |
|  | 70 | Male |  |
|  | 63 | Male |  |
|  | 73 | Female |  |

**Supplementary Table 2.**
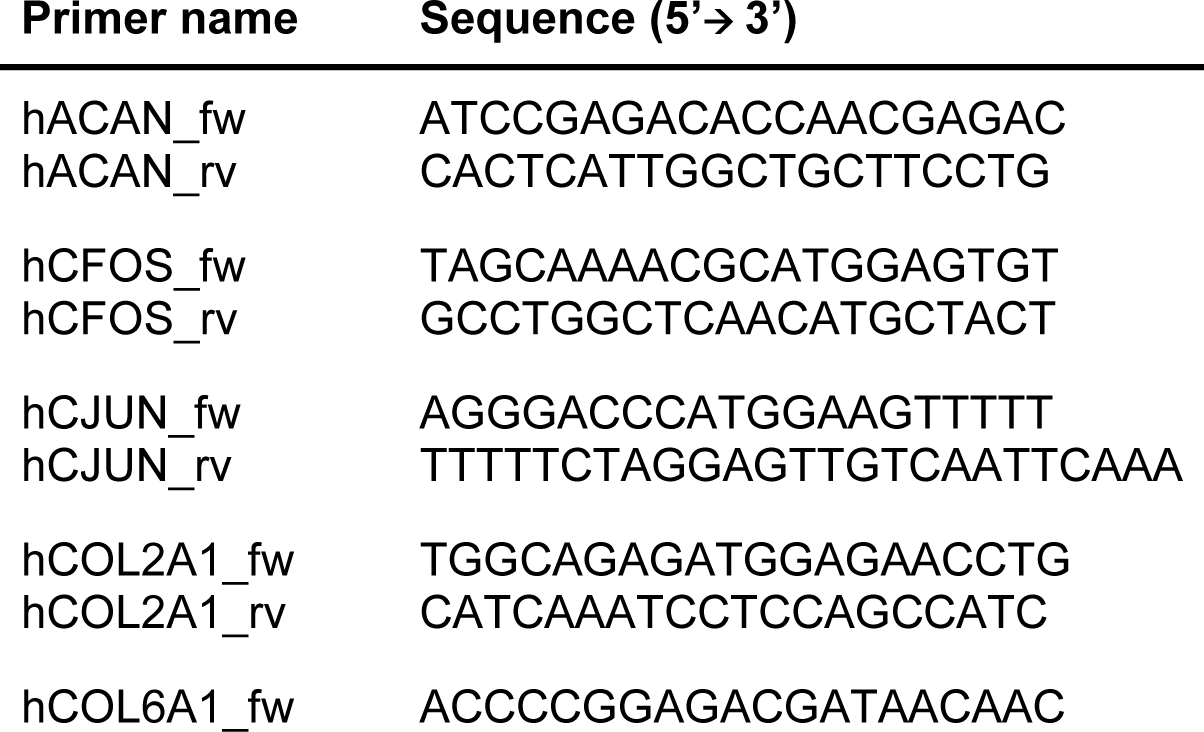
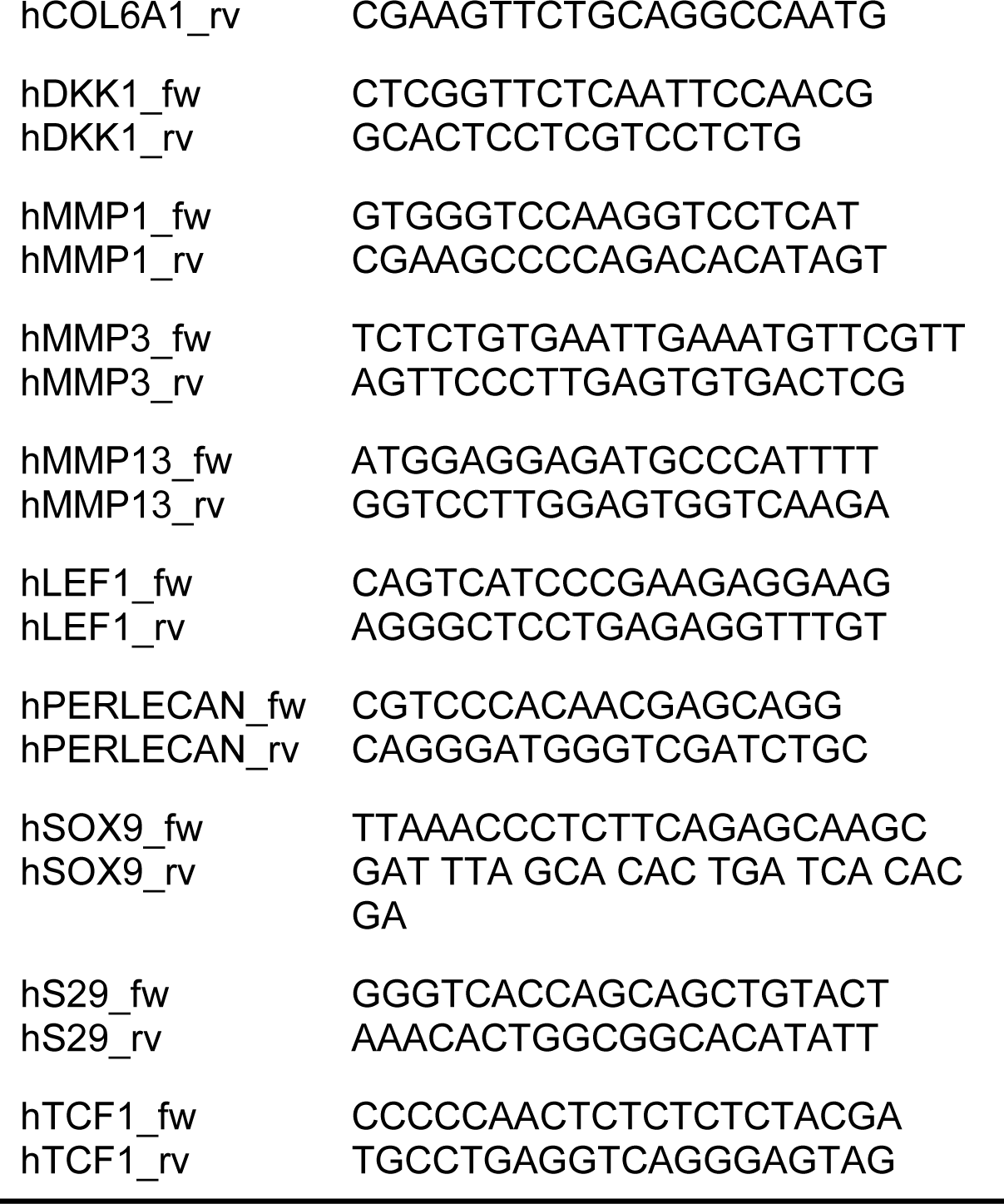
Human primers used in qPCR.

**Supplementary Table 3.** Statistical analysis details.

| **Figure 2B** | Linear mixed model | t value | p value |
| --- | --- | --- | --- |
| *TCF-1* | *CHIR* | 3.167 | 0.00380 |
|  | *LOAD* | 3.089 | 0.00461 |
|  | *INTERACTION* | -1.750 | 0.09156 |
| *cJUN* | *CHIR* | -0.168 | 0.867820 |
|  | *LOAD* | 4.500 | 0.000117 |
|  | *INTERACTION* | -0.396 | 0.695359 |
| *SOX9* | *CHIR* | -2.768 | 0.01007 |
|  | *LOAD* | 3.084 | 0.00468 |
|  | *INTERACTION* | -0.240 | 0.81182 |
| *COL2A1* | *CHIR* | -3.481 | 0.00171 |
|  | *LOAD* | 0.109 | 0.91426 |
|  | *INTERACTION* | 0.303 | 0.75656 |
| *ACAN* | *CHIR* | -3.466 | 0.00178 |
|  | *LOAD* | -1.319 | 0.19824 |
|  | *INTERACTION* | 0.052 | 0.95916 |
| *COL6* | *CHIR* | -0.711 | 0.483 |
|  | *LOAD* | 0.548 | 0.588 |
|  | *INTERACTION* | -0.988 | 0.332 |
| *Perlecan* | *CHIR* | -0.987 | 0.33250 |
|  | *LOAD* | 2.973 | 0.00614 |
|  | *INTERACTION* | -0.845 | 0.40542 |
| *MMP1* | *CHIR* | -0.357 | 0.72453 |
|  | *LOAD* | 4.402 | 0.00019 |
|  | *INTERACTION* | -1.022 | 0.31685 |
| *MMP3* | *CHIR* | -0.297 | 0.768 |
|  | *LOAD* | 1.674 | 0.104 |
|  | *INTERACTION* | 0.072 | 0.943 |
| *MMP13* | *CHIR* | 1.819 | 0.0801 |
|  | *LOAD* | 4.926 | 3.71e-05 |
|  | *INTERACTION* | -2.164 | 0.0395 |

## 13. Linear Mix Model R script

\## Linear mixed model for interpreting gene expression upon loading in CHIR treated Non-OA patients

\## Preparation of R

library(car) # regression tools

library(rstatix) # pipe friendly data analysis

library(tidyverse) # facilitate data processing including ggplot2 graphics

library(ggpubr) # additional graph features

library(lme4) # linear mixed models

library(emmeans) # estimates and confidence intervals

library(MASS) #distribution

library(multcomp)

library(lmerTest)

library(EnvStats)

library(readr)

library(effects)

library(sjPlot)

library(Matrix)

library (lattice)

\## Preparing the data

dataset <-read_csv(“datarocio_FoldChange.csv”)

dataset$ID <-factor(dataset$ID) # set as factor

dataset$CHIR <-factor(dataset$CHIR) # set as factor

dataset$Loading <-factor(dataset$Loading) # set as factor

dataset2 <-dataset # define mirror data set for transformation

dataset2[,4:ncol(dataset2)] <-log10(dataset2[,4:ncol(dataset2)])# log transform gene expression values

dataset2[,4:ncol(dataset2)] <-apply(dataset2[,4:ncol(dataset2)], 2, as.numeric)

\## Loop for all variables

for(variable in names(dataset2[,4:ncol(dataset2)])){

lmm_gene <-lmer(get(variable) ∼ CHIR * Loading + (1 | ID), REML = FALSE, data = dataset2)

qqplot_model <-qqmath(lmm_gene, id=0.05)

assumption_plots <-plot_model(lmm_gene, type=’diag’)

Summary_lmm <-summary(lmm_gene)

post_hoc_comparison <-emmeans(lmm_gene, list(pairwise ∼ CHIR*Loading), adjust = “tukey”, infer = TRUE)

cat(“Gene:”, variable, “\n”)

cat(“Checking normality:”, variable, “\n”)

print(qqplot_model)

cat(“Checking Assumptions for:”, variable, “\n”)

print(assumption_plots)

cat(“Linear Mixed Model for:”, variable, “\n”)

print(Summary_lmm)

cat(“Post Hoc comparison of”, variable, “\n”)

print(post_hoc_comparison)

}

## References

1. Walter SG, Ossendorff R, Schildberg FA. Articular cartilage regeneration and tissue engineering models: a systematic review. Arch Orthop Trauma Surg. 2019;139(3):305–316. doi:10.1007/s00402-018-3057-z

2. Sophia Fox AJ, Bedi A, Rodeo SA. The Basic Science of Articular Cartilage: Structure, Composition, and Function. Sports Health: A Multidisciplinary Approach. 2009;1(6):461–468. doi:10.1177/1941738109350438

3. Knobloch TJ, Madhavan S, Nam J, Agarwal Jr, S, Agarwal S. Regulation of Chondrocytic Gene Expression by Biomechanical Signals. Crit Rev Eukaryot Gene Expr. 2008;18(2):139–150. doi:10.1615/CritRevEukarGeneExpr.v18.i2.30

4. Patel JM, Wise BC, Bonnevie ED, Mauck RL. A Systematic Review and Guide to Mechanical Testing for Articular Cartilage Tissue Engineering. Tissue Eng Part C Methods. 2019;25(10):593–608. doi:10.1089/ten.tec.2019.0116

5. Zhao Z, Li Y, Wang M, Zhao S, Zhao Z, Fang J. Mechanotransduction pathways in the regulation of cartilage chondrocyte homoeostasis. J Cell Mol Med. 2020;24(10):5408–5419. doi:10.1111/jcmm.15204

6. Gilbert SJ, Blain EJ. Cartilage mechanobiology: How chondrocytes respond to mechanical load. In: Mechanobiology in Health and Disease. Elsevier; 2018:99–126. doi:10.1016/B978-0-12-812952-4.00004-0

7. Global burden of disease study 2019 (GBD 2019) results. Published online 2020.

8. Hunter DJ, Bierma-Zeinstra S. Osteoarthritis. The Lancet. 2019;393(10182):1745-1759. doi:10.1016/S0140-6736(19)30417-9

9. Kitagaki J, Iwamoto M, Liu JG, Tamamura Y, Pacifci M, Enomoto-Iwamoto M. Activation of β–catenin-LEF/TCF signal pathway in chondrocytes stimulates ectopic endochondral ossification. Osteoarthritis Cartilage. 2003;11(1):36–43. doi:10.1053/joca.2002.0863

10. Dong YF, Soung DY, Schwarz EM, O’Keefe RJ, Drissi H. Wnt induction of chondrocyte hypertrophy through the Runx2 transcription factor. J Cell Physiol. 2006;208(1):77–86. doi:10.1002/jcp.20656

11. Monteagudo S, Lories RJ. Cushioning the cartilage: a canonical Wnt restricting matter. Nat Rev Rheumatol. 2017;13(11):670–681. doi:10.1038/nrrheum.2017.171

12. Lories RJ, Monteagudo S. Review Article: Is Wnt Signaling an Attractive Target for the Treatment of Osteoarthritis? Rheumatol Ther. 2020;7(2):259–270. doi:10.1007/s40744-020-00205-8

13. Dell’Accio F, De Bari C, El Tawil NM, Barone F, Mitsiadis TA, O’Dowd, et al. Activation of WNT and BMP signaling in adult human articular cartilage following mechanical injury. Arthritis Res Ther. 2006;8(5):R139. doi:10.1186/ar2029

14. Huang G, Chubinskaya S, Liao W, Loeser RF. Wnt5a induces catabolic signaling and matrix metalloproteinase production in human articular chondrocytes. Osteoarthritis Cartilage. 2017;25(9):1505–1515. doi:10.1016/j.joca.2017.05.018

15. Wang X, Cornelis FMF, Lories RJ, Monteagudo S. Exostosin-1 enhances canonical Wnt signaling activity during chondrogenic differentiation. Osteoarthritis Cartilage. 2019;27(11):1702–1710. doi:10.1016/j.joca.2019.07.007

16. Timmermans RGM, Bloks NGC, Tuerlings M, van Hoolwerff M, Nelissen RGHH, van der Wal RJP, et al. A human in vitro 3D neo-cartilage model to explore the response of OA risk genes to hyper-physiological mechanical stress. Osteoarthr Cartil Open. 2022;4(1):100231. doi:10.1016/j.ocarto.2021.100231

17. Occhetta P, Mainardi A, Votta E, Vallmajo-Martin Q, Ehrbar M, Martin I, et al. Hyperphysiological compression of articular cartilage induces an osteoarthritic phenotype in a cartilage-on-a-chip model. Nat Biomed Eng. 2019;3(7):545–557. doi:10.1038/s41551-019-0406-3

18. Praxenthaler H, Krämer E, Weisser M, Hecht N, Fischer J, Grossner T, et al. Extracellular matrix content and WNT/β-catenin levels of cartilage determine the chondrocyte response to compressive load. Biochimica et Biophysica Acta (BBA) - Molecular Basis of Disease. 2018;1864(3):851–859. doi:10.1016/j.bbadis.2017.12.024

19. van Caam A, Cocca RA, Farran AJ, Mohanraj B, Meloni G, Mauck R, et al. Perlecan expression is strongly reduced in aging cartilage but increased by physiological loading. Osteoarthritis Cartilage. 2014;22:S60. doi:10.1016/j.joca.2014.02.124

20. Blain EJ, Mason DJ, Duance VC. The effect of cyclical compressive loading on gene expression in articular cartilage. Biorheology. 2003;40(1-3):111–117.

21. Parks WC, Wilson CL, López-Boado YS. Matrix metalloproteinases as modulators of inflammation and innate immunity. Nat Rev Immunol. 2004;4(8):617–629. doi:10.1038/nri1418

22. Rose BJ, Kooyman DL. A Tale of Two Joints: The Role of Matrix Metalloproteases in Cartilage Biology. Dis Markers. 2016;2016:1–7. doi:10.1155/2016/4895050

23. Freitas-Rodríguez S, Folgueras AR, López-Otín C. The role of matrix metalloproteinases in aging: Tissue remodeling and beyond. Biochimica et Biophysica Acta (BBA) - Molecular Cell Research. 2017;1864(11):2015–2025. doi:10.1016/j.bbamcr.2017.05.007

24. Martel-Pelletier J, Barr AJ, Cicuttini FM, Conaghan PG, Cooper C, Goldring MB, et al. Osteoarthritis. Nat Rev Dis Primers. 2016;2(1):16072. doi:10.1038/nrdp.2016.72

25. Dolzani P, Assirelli E, Pulsatelli L, Meliconi R, Mariani E, Neri S. Ex vivo physiological compression of human osteoarthritis cartilage modulates cellular and matrix components. PLoS One. 2019;14(9):e0222947. doi:10.1371/journal.pone.0222947

26. Goldring MB, Otero M. Inflammation in osteoarthritis. Curr Opin Rheumatol. 2011;23(5):471-478. doi:10.1097/BOR.0b013e328349c2b1

27. Poole AR. Osteoarthritis as a Whole Joint Disease. HSS Journal. 2012;8(1):4–6. doi:10.1007/s11420-011-9248-6

28. Grumolato L, Liu G, Mong P, Mudbhary R, Biswas R, Arroyave R, et al. Canonical and noncanonical Wnts use a common mechanism to activate completely unrelated coreceptors. Genes Dev. 2010;24(22):2517–2530. doi:10.1101/gad.1957710

29. Nalesso G, Sherwood J, Bertrand J, Pap T, Ramachandran M, De Bari C, et al. WNT-3A modulates articular chondrocyte phenotype by activating both canonical and noncanonical pathways. Journal of Cell Biology. 2011;193(3):551–564. doi:10.1083/jcb.201011051

30. Dantas LO, Salvini T de F, McAlindon TE. Knee osteoarthritis: key treatments and implications for physical therapy. Braz J Phys Ther. 2021;25(2):135–146. doi:10.1016/j.bjpt.2020.08.004

31. Skou ST, Roos EM. Physical therapy for patients with knee and hip osteoarthritis: supervised, active treatment is current best practice. Clin Exp Rheumatol. 2019;37 Suppl 120(5):112–117.

32. Kraus VB, Blanco FJ, Englund M, Karsdal MA, Lohmander LS. Call for standardized definitions of osteoarthritis and risk stratification for clinical trials and clinical use. Osteoarthritis Cartilage. 2015;23(8):1233–1241. doi:10.1016/j.joca.2015.03.036

